# Raising the Connectome: the emergence of neuronal activity and behavior in *Caenorhabditis elegans*

**DOI:** 10.1101/2020.01.06.896308

**Authors:** Bradly Alicea

## Abstract

The differentiation of neurons and formation of connections between cells is the basis of both the adult phenotype and behaviors tied to cognition, perception, reproduction, and survival. Such behaviors are associated with local (circuits) and global (connectome) brain networks. A solid understanding of how these networks emerge is critical. This opinion piece features a guided tour of early developmental events in the emerging connectome, which is crucial to a new view on the connectogenetic process. Connectogenesis includes associating cell identities with broader functional and developmental relationships. During this process, the transition from developmental cells to terminally differentiated cells is defined by an accumulation of traits that ultimately results in neuronal-driven behavior. The well-characterized developmental and cell biology of *C. elegans* will be used to build a synthesis of developmental events that result in a functioning connectome. Specifically, our view of connectogenesis enables a first-mover model of synaptic connectivity to be demonstrated using data representing larval synaptogenesis. In a first-mover model of Stackleberg competition, potential pre- and postsynaptic relationships are shown to yield various strategies for establishing various types of synaptic connections. By comparing these results to what is known regarding principles for establishing complex network connectivity, these strategies are generalizable to other species and developmental systems. In conclusion, we will discuss the broader implications of this approach, as what is presented here informs an understanding of behavioral emergence and the ability to simulate related biological phenomena.

**Highlights:**

- we can understand the complexity of connectomes in terms of their emergence from embryogenetic precursors, connection dynamics, and relationship to organismal behavior.
- a first-mover competition model can explain how neuronal cells follow a specific set of heuristic strategies to form chemical synapses with other cells in larval development.
- the timing and relative order of terminal differentiation in *Caenorhabditis elegans* are shown to have both subtle and consequential effects on patterns of connectivity.
- a correspondence is established between the emergence of small connectomes and the emergence of specific behavioral outcomes in both animal and *in silico* models.

## Introduction

The field of connectomics provides opportunities to view the structure and function of nervous systems in a new light. The proliferation of data for small connectomes such as those found in the nematode *Caenorhabditis elegans* allows us to study the connected nervous system at both the single cell and systems level (Berck et.al, 2016; Eichler et.al, 2017; Ko et.al, 2013). Emerging functional perspectives on genomic contributions (Barabasi and Barabasi, 2020) and statistical regularities (Takagi, 2017) of the connectome allow for a flexible, multiscale model of biology and behavior. In this opinion piece, these trends in the study of connectomics are leveraged to identify associations between the various properties and behaviors of terminally differentiated neurons and their presence at various points in embryogenesis. An informatics approach is employed to unify the individual steps of connectogenesis and reveal the systemic regularities that result in a functionally unified network. The term *connectogenesis* is proposed as a means to describe the process of nervous system development in the greater context of cellular interactions, behavioral function, and the organismal life cycle.

As we utilize multiple sources of data, a number of assumptions are made that might obfuscate the differences between various stages in life-history. For example, it is important to distinguish between the timing of events associated with the adult state of cells (such as gene expression) and the embryonic state of cells with the same identity (Hobert et.al, 2016). Cells emerge in a far different physiological milieu than those of the adult connectome. One counter-argument to our approach involves new differentiated neurons in development that do not have identical gene expression state to that found in adult versions of these same cells. *C. elegans* genome encodes a multitude of proteins that contribute to neuronal activity and communication, particularly across the various states of development. On the other hand, cellular state tends to be conserved when comparing development and adulthood. Identifying genes expressed in the adult versions of the cells might be in fact useful indicators of their developmental origins and the effects of as of yet unidentified developmental constraints. Based on these partial patterns of adult connectivity, generalizations are made with respect to the structural and functional features of a developing connectome (Kaiser, 2017).

To more fully understand the sequential assembly aspect of connectogenesis, we introduce a first-mover model of cell-cell competition for establishing synaptic connectivity. A dataset of synaptic establishment and dynamics over the course of development (Witvliet et.al, 2020) is used to uncover strategies favored by individual cells in the course of establishing presynaptic and postsynaptic relationships. First-mover dynamics are based on the principle of Stackleberg competition (Simaan & Cruz, 1973), which is a competitive model of sequential actions. In the developmental biological context, first-movers have the advantage of not being directly subject to developmental contingencies (Morton, 1986). A first-mover model is meant to capture the process of growth, differentiation, and associated interactions defining the transition from emerging structure to organized function (Vogelstein et.al, 2019). The first-mover model of Stackleberg competition was first applied to developmental systems in Stone et.al (2018), and is used here to represent the evolution of interactions between the emergence of newly-born neurons, asymmetric ion channel expression amongst bilateral pairs, and the formation of neural circuits.

A parallel area of inquiry involves exactly when and under what conditions these terminally differentiated neurons begin to produce organismal-level behaviors. It has been suggested that structural features emerge earlier in development than functional features (Cao et.al, 2016). In general, the first neuronal cells emerge well before the first synaptic connections between cells are established. In a related fashion, the first goal-directed behaviors require synaptic connections to be established between neuronal and muscle cells. Our contribution to this understanding is to flesh out the structure of this timing relative to differentiation events. This includes making connections between the terminal differentiation of specific neurons in development, their eventual functional identity, and the systems-level significance of this temporal process. These relationships help us understand the tempo and scope of connectogenesis relative to the adult phenotype.

In this paper, we provide a guided tour of early developmental events of *C. elegans* connectome. Such a perspective is lacking in the *C. elegans* literature, and is broadly applicable to the study of developmental dynamics. We build on the work of Alicea (2018), where the construction of partial connectomes for multiple time points in embryonic development demonstrates how the connectome emerges from developmental cells. Using information from multiple sources, we build a timeline that spans from the birth of the first neuron to the emergence of the first behaviors (Figure 1). We will also ascribe strategies to individual cells that enable them to form synaptic connections. This information processing capacity is the product of cellular identity, which is a combination of relative factors (anatomical position or birth order) and internal factors (molecular properties). While we do not identify causal factors, they might be better understood using simulation approaches included in the discussion section.

**Figure 1.**
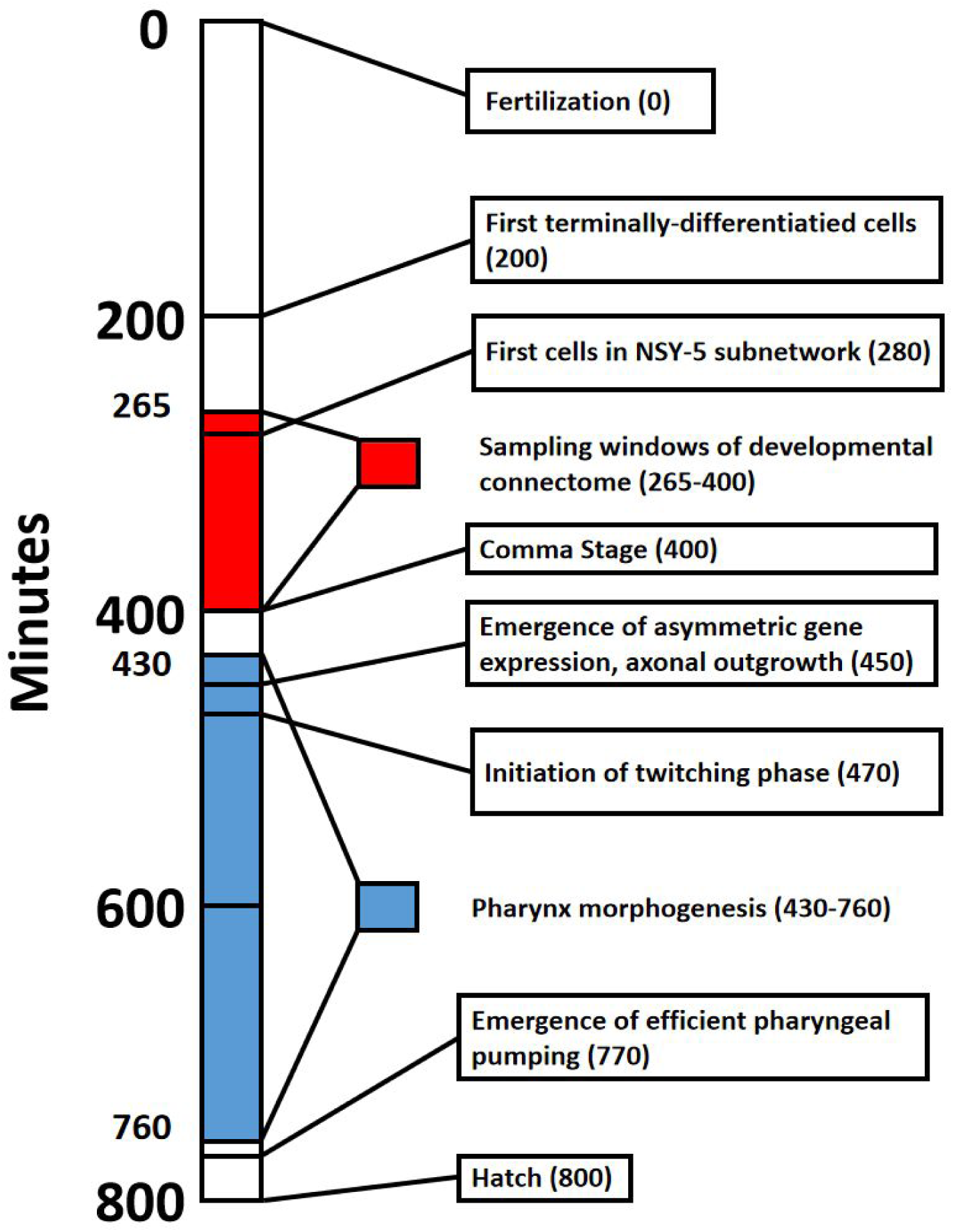
Timeline that shows major events in *C. elegans* embryogenesis from fertilization (0 minutes) to hatch from the egg (800 minutes). Red shading denotes the sampling window for cells of the developmental connectome, spanning from 265 to 400 minutes post-fertilization. Blue shading denotes Pharynx morphogenesis, which was selected to highlight the formation of a functional trait in the emerging adult phenotype.

### Establishing the Timing of Connectogenetic Events

The nematode *C. elegans* has been chosen for this study for two reasons. The first is a well-established community of open resources that enable the integration of connectomic, developmental, and molecular data. Secondly, the straightforward mapping between developmental cell lineages, differentiating neurons in the embryo, and the adult phenotype helps us to establish associations between the two stages of life-history. While *C. elegans* neurodevelopment is deterministic, the connectome is both small and functionally diverse. The adult connectome consists of 302 neurons in the hermaphrodite (White et.al, 1986) and 385 cells in the male (Cook et.al, 2019). We can thus trace the individual components of connectogenesis in a tractable manner. The lack of functional ambiguity in our model organism provides both well-established patterns of connection and well-defined structural features (Varshney et.al, 2011).

Characterization of differentiation events of the connectome are predicted from observations of embryo phenotypes. In this paper, an informatics approach is used to associate identified terminally differentiated neuronal cells with their adult identities. Yet we must recognize that the expression of genes and ion channels are not synonymous with the first appearance of a specific cell. Even though *C. elegans* has a deterministic developmental trajectory with respect to cell lineage and identity, this process involves the gradual expression and maintenance of transcription factors. The terminal selector (Hobert, 2016a), transcriptional mechanism serves to simultaneously initiate and maintain cell fate choice. Protectors/stabilizers (Zheng and Chalfie, 2016) then ensure the smooth, uninterrupted expression of terminal selectors over time. The notion that neuronal differentiation is a complex process that unfolds over time is most important in understanding connectogenesis.

It is imperative that details regarding the so-called neuronal identity of these newly-integrating cells be understood in order to understand connectogenesis while also relating this process to functional and behavioral phenomena (Hobert, 2016b). *C. elegans* neurons undergo a multipotency-to-commitment transition (MCT), in which neuronal cells are born from a diverse developmental cell lineage that also gives rise to muscle and epithelial cells (Rothman and Jarriault, 2019). This may provide a basis for great electrophysiological diversity across cells in very distinct subcircuits relevant to behavior. For example, there is a difference between functional asymmetry and cell lineage asymmetry which is partially driven by the expression and function of ion channels. While the anatomical distribution of neurons generally remains conserved throughout development (Nicosia et.al, 2013; Alicea, 2017), a number of post 400-minute transformations define connectivity and ultimately organismal-level behaviors. These transformations are often connected to the increasing asymmetry in phenotype as the spherical egg becomes an asymmetrical organism (Gordon and Gordon, 2016).

### First-mover Model of Connectogenesis

A first-mover model of developmental emergence is used to tie together many of these developmental processes. This has parallels with the game-theoretic model of Khajezade et.al (2019), in which utility functions derived from a specialized network are used to model optimal connectivity in the frontal network of *C. elegans* connectome. However, their model did not assume that development is a generative process (from a single cell to a complex phenotype). Our results propose a developmental-specific competitive mechanism (first-mover competition), a formal analysis of observed data (establishment of synaptogenesis), and generalizing the results to a systems-level context.

The first-mover model describes neurons that are born and eventually connect to an expanding network. As a sequential process, pairs of neurons engage in a leader-follower dynamic to establish connections. In this type of competition, there are first movers (leaders), who establish the nature of the dyadic relationship, as well as second movers (followers), who are constrained by the outcome of the first move. In this instance, a single move is a connection between two cells, initiated by the first mover. In a sequential interaction, initial two-player competitions and their associated outcomes provide information but also increasingly limited options for later competitions. This might lead to suboptimal outcomes or the selection of multiple strategies for a single cell identity.

In recent work on developmental Braitenberg Vehicles (dBVs in Dvoretskii et.al, 2020), we can see how the first-mover model works. Braitenberg Vehicles (Braitenberg, 1984) are simple models that approximate simple organisms that behave in response to simple environmental cues. In the developmental version, we grow a nervous system from a simple sensor-effector mapping to a complex network of intermediate cells and connections. In this case, cells are introduced and remain uncoupled until an analogy to synaptic connections are established. As new cells are introduced into the nervous system, they must connect to existing members of the network. From this initial condition, it becomes obvious that each cell uses some sort of criterion to attach to the network, while cells that enter the network at a later point in time are constrained in terms of both entry position and possible partners. In real organisms, chemical cues and genetic mechanisms govern supporting factors such as axonogenesis and intercellular signalling. Yet this network scaffolding can only act to enable connectivity, not impose temporal order on the expanding connectome. The dBV model demonstrates how temporal order proceeds during connectogenesis, and results in a hypothesis: cells that are established (or differentiated) first will also wire to the network first, and occupy prime structural and functional positions in the network.

We can look to real neural systems to confirm some of the intuitions raised by the toy model. For example, so-called pioneering axons determine the later pathways for follower axons that occur later in development (Chedotal and Richards, 2010). Furthermore, as developmental plasticity proceeds post-birth, cells that connect first serve to establish the experience-dependent properties of the network (Tierney and Nelson, 2009). This might include the emergence of processing centers or selective enhancement of certain pathways over others. Kim and Kaiser (2014) even suggest that in *C. elegans* connectome, the establishment of functional connectivity proceeds as a means to maximize the distribution of local information as well as a means to lessen algorithmic entropy. Both of these outcomes suggest that while birth order might be important in establishing temporal order in an expanding connectome, it does not always act in a deterministic manner. In other words, while this criterion is generally explanatory, it is not a universal principle. Nor is birth time preference immediately apparent in the data; each cell establishes multiple connections to the connectome as a whole, and these connections occur at different points in time. Some connections are more constrained by their relative timing than others made by the same cell. Nevertheless, we can assess the presence of average behaviors to capture this principle of order, in addition to ascribing strategies to these tendencies.

During the course of connectogenesis, a single cell establishes chemical connections with multiple pre- and postsynaptic partners. In establishing these connections amongst a population of neuronal cells, each cell exhibits a strategy based on maturity from birth time. Older cells are cells that are born first (earlier in embryogenesis), while newer cells are born second (later in embryogenesis). We measure these tendencies in terms of average behavior. Based on this heuristic, an individual cell can exhibit strategies that vary in a number of ways. Some strategies will restrict the cell to be totally committed to a particular strategy. One example of this would involve our cell only serving as a presynaptic partner to younger cells (born second) and postsynaptic partner to older cells (born first). Alternatively, a strategy might be defined by a particular type of bias, such as only excluding postsynaptic partnership with younger cells. Other strategies might simply exhibit a particular tendency over time, preferring older postsynaptic partners but not choosing them exclusively.

The goal of a first mover advantage and the shifting dynamics that new agents with heterogeneous strategies provides a means to achieve leader-follower behavior. This in turn imposes assortative order upon this developmental network. Our first-mover model provides us with a hypothetical model for the structure of developmental wiring. An example of this is the fitness-based connectivity criterion of Bianconi and Barabasi (2001). Given a selection scheme such as rank or tournament selection (Hamblin, 2012), neurons connect at different rates to its neighbors based on some functional objective.

### Outline of Approach

Our analysis proceeds in four parts. The first part involves an analysis of cells at different points in time (Alicea et.al, 2018), allowing for an inventory of the various tissue and organ types represented by newly differentiated neurons. Ion channel data collected from WormBase (Harris et.al, 2019) are used to identify key genes, associated cells, and annotations. Determining the relationships between functional gene identities, specific ion channels, and functional annotations will be enhanced by using data sets (see Methods) developed at the OpenWorm Foundation (Sarma et.al, 2018). This analysis yields information regarding ion channel types present at a specific time point in embryogenesis.

In terms of the latter, the data set featured in (Packer et.al, 2019) is used in conjunction with annotation metadata and embryonic gene expression information to draw conclusions regarding the phylogenetic and developmental origins of the connectome. Of particular interest is the link between ion channels expressed by neuronal cells present in the embryo and the lineage-specific expression of ion channel-related and other associated genes. The final component relies upon how stereotypical *C. elegans* behaviors are heavily influenced by genetics to establish a link between the presence of certain neurons and the potential for early forms of chemosensation, thermosensation, and even learning and memory (Walker et.al, 2019).

## Methods

To make associations between the identity of new neurons and their later molecular identity, an informatics approach is utilized. Data from the OpenWorm Foundation’s easily-accessible collection of terminally-differentiated cell birth times, cell names, and annotations are required to reconstruct the developmental connectome. Verification of cell birth time is done using the accessible and interactive resource of Bhatla (2011). Additional annotation of each cell and where they fit into what we know about the *C. elegans* nervous system have been retrieved from WormBase, which serves as an accessible community resource. References from the scientific literature are brought to bear for providing context to cells that emerge in development. The WormBase and literature review resources are especially useful in constructing a timeline of developmental events shown in Figure 1. The synaptic data analysis from Witvliet et.al (2020) demonstrates how these cells make synaptic connections once the egg has hatched.

### OpenWorm Resources

Functional annotations are courtesy of Stephen Larson and Mark Watts from datasets provided by the OpenWorm Foundation (http://openworm.org). The DevoWorm group has also assembled a number of bioinformatic conversions that have been used to track the birth times and developmental lineage of each cell in the *C. elegans* connectome. These data can be found on Github (https://github.com/devoworm/DevoWorm) and the Open Science Framework (Alicea, 2018).

### WormBase

Supplemental information about protein expression and cells of interest are retrieved from WormBase Version:WS273 (https://www.wormbase.org/). WormBase (Harris et.al, 2019) is a *C. elegans* community resource that allows us to map between specific cells and the properties of their adult phenotypes.

### Definitions of cell types

Developmental cells are part of an actively-dividing invariant cell lineage in which fate acquisition is deterministic, while terminally differentiated cells are those which have both reached a terminal division and have acquired a terminal functional state. While postembryonic plasticity may result in variability in a cell’s full identity across life-history (Hobert et.al, 2016), terminally differentiated cells observed in the embryo or larval worm are not identical to their adult counterparts.

### Definition of NSY-5 network

The NSY-5 network is a gap junction network consisting of 32 cells (Chuang et.al, 2007). There are 28 left/right pairs of cells in this network: ADA, ADF, ADL, AFD, AIM, AIZ, ASH, ASI, ASK, AWB, AWC, BAG, and RIC. As opposed to our time-threshold networks (e.g. connectogenesis at 300 minutes post-fertilization), neurons included in this network are defined by their expression of NSY-5 (Chuang et.al, 2007).

### Definition of synaptic states

The synapse classifications of Witvliet et.al (2020) are based on the assumption that all synaptic connections in the adult connectome are hard-wired (or stereotyped). During the last stages of connectogenesis (larval development), patterns of connectivity can deviate from this assumption. Transient synapses (43% of synaptic connections in the adult connectome) are highly-variable synapses that are not always present across individual worms across the C. elegans species. There is also no clear trend in synapse number across life-history. Developmental synapses (14% of synaptic connections in the adult connectome) are dynamic across development, and reflect the formation of new connections and elimination of superfluous connections. Stable synapses (43% of synaptic connections in the adult connectome) are stereotyped connections present from hatch until the end of the L4 stage, as the relative strength of pre- and postsynaptic cells are also maintained over this period (Witvliet et.al, 2020).

### Data for first-mover analysis

2,809 potential cell-cell pairwise (*i,j*) relationships denoting pre and postsynaptic relationships are used to determine the first-mover behavior of 58 cells in the adult connectome are taken from Witvliet et.al (2020). To determine which cells are more likely to be first and second movers, a measure called *birth time difference* is calculated. Birth time difference measures the difference between the birth time of a single presynaptic neuron and the birth time of a single postsynaptic neuron. Negative values represent presynaptic cells that were born first, while positive values represent presynaptic cells that were born second. In calculating times for our strategies, the mean time for all pairwise comparisons for a single row or column in the pairwise matrix i,j are used. These times are further decomposed into positive and negative components, which are then used in calculating the birth time difference. All notes, figures, and code associated with the first-mover model are located in a Github repository (https://github.com/devoworm/Raising-the-Worm-Brain).

Each neuron has a suite of strategies available at any given time. Such profiles are heterogeneous, and can influence the gene expression of newly-connected cells based on the principle of first-mover. For the example shown in Table 1, the cell in question will act as both a pre- and postsynaptic partner. For a majority of connections, the cell in question will establish connectivity with a postsynaptic partner born later and a presynaptic partner born earlier (75% of the time). Occasionally, this heuristic will be violated, and some strategies might be more specific in excluding presynaptic partners born later and/or postsynaptic partners born earlier than our hypothetical cell (100% of the time). In such a case, the one or more of the combinations in Table 1 will occur with a frequency of zero (0% of the time).

**Table 1.** Probabilistic relationships between synaptic partners and birth times. Each cell in the connectome will exhibit a proportion of these four relationships, which in turn determines the strategy exhibited. Note that in this example, the presynaptic, born first and postsynaptic, born second relationships are linked, so that the percentages are derived by summing diagonally.

|  | Born First | Born Second |
| --- | --- | --- |
| Presynaptic | 0.375 | 0.125 |
| Postsynaptic | 0.125 | 0.375 |

### Definition of observed strategies

By separating the developmental birth time data into positive and negative components for both presynaptic and postsynaptic partners, five strategies have been identified. This includes four strategies that involve some form of negative-positive (N-P) coupling, and one strategy that involves positive-negative (P-N) coupling. A graphical representation of these strategies in terms of relationships between connectivity and birth times are shown in Figure 2.

**Figure 2.**
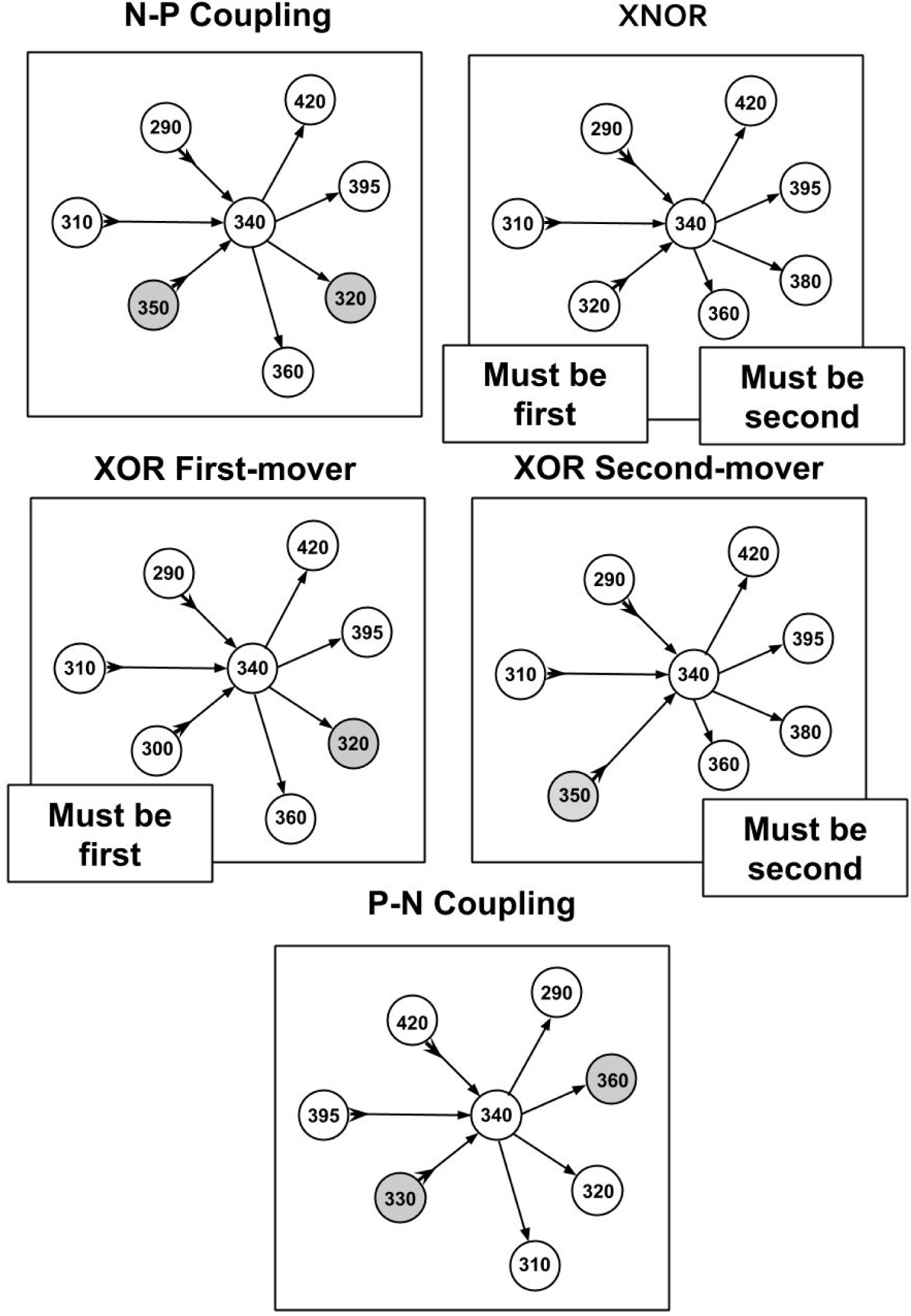
Diagram of strategies and relationships between cell making connections and cell times of synaptic partners. Presynaptic relationships are denoted with a forward arrow → towards the postsynaptic partner while a postsynaptic relationship is denoted with an arrow towards the cell making connections that also has a tail at the reverse end ↣. Gray circles (nodes) are cells that violate the primary strategy (but make up a minority of incoming and outgoing connections overall).

1. Negative-Positive (N-P) Coupling. For our sample of synaptic formation data, this strategy is employed for roughly 40-50% of connections. N-P coupling includes three other strategies which are more focused aspects of the synaptic pairwise relationship: XOR First Mover, XOR Second Mover, and XNOR. When a cell exhibits the N-P coupling strategy, the presynaptic cell is born earlier in developmental time on average than its postsynaptic counterpart.
2. Exclusive OR (XOR) First Mover. This strategy is employed for around 7-8% of connections, and can include both pre- and post-synaptic partners born first. The XOR first-mover strategy is a refinement of N-P in that it excludes presynaptic partners born after the cell in question.
3. Exclusive OR (XOR) Second Mover. This mixed strategy is employed for around 20% of connections. This strategy can include both pre- and postsynaptic partners born second. XOR second-mover only excludes postsynaptic partners born first.
4. Exclusive NOR (XNOR). In our sample, this strategy is exhibited around 20% of connections. As an even more exclusive version of the XOR strategies, XNOR strategies include only those connections where the presynaptic cell is born first and the postsynaptic cell is born second.
5. The Positive-Negative (P-N) coupling category contains cells that have the opposite relationship between pre- and postsynaptic cells relative to N-P coupling. For cells deploying this strategy, on average the presynaptic cells are born later than their postsynaptic partner. P-N coupling is a pure strategy which does not overlap with any other strategies.

### Utilities for first-mover model

Employing a series of strategies over time results in a payoff for each cell that is semi-independent of its specific set of preferred strategies. The payoffs determine the costs and benefits of what it means to be a first-mover, a subsequent-mover, or a random mover. Our estimated components of utility can be defined as follows: *FREQ* represents empirically-observed frequencies in development, *M* represents an estimate of the molecular milieu), and *N* represents an estimate of the neurophysiological milieu. This can be estimated in a context-dependent manner, while the formulation is shown for the *C. elegans* connectome in Table 2.

**Table 2.** Payoff matrix for a presynaptic cell that makes a connection with another cell in the nervous system.

|  | **First-mover** | **Subsequent-Mover** | **Random** |
| --- | --- | --- | --- |
| **First-mover** | - | $FREQ * (M + N)$ | $FREQ * 0.5$ |
| **Subsequent-Mover** | $1 - FREQ * (M + N)$ | - | $FREQ / 0.5$ |
| **Random** | $FREQ * 0.5$ | $FREQ / 0.5$ | - |

## Results

The first part of this analysis will proceed by walking through the progression of developmental events in two orthogonal ways. One characterization involves looking at temporal progression. Our inquiry begins before the differentiation of neuronal cells, and the proto-connectome as it exists at five points during embryogenesis (265, 280, 290, 300, and 400 minutes post-fertilization). These networks are defined in Alicea (2018), where the progression of connectogenesis is defined indirectly in terms of cells present.

The second characterization involves characterizing molecular properties and events associated with the terminally differentiated neurons. One example of this is the NSY-5 subnetwork (see Methods), whose constituents are defined by their molecular properties (Chuang et.al, 2007). The NSY-5 network demonstrates how developmental asymmetry begins to emerge in connectogenesis. While this asymmetry is limited to the exchange of small molecules between its subnetwork constituents, it also allows us to think about the adult connectome as a developmentally contingent structure. Yet there may also be consequences of NSY-5 subnetwork connectogenesis on the emergence of behavior, as *nsy-5/inx-19*^*-*^ mutants that lack a functioning AWC cell pair exhibit deficits in electrotactic and associated navigation behaviors (Chrisman et.al, 2016). Additionally, as the cholinergic network emerges in development, so do the roots of functional criticality (highly-connected cells that are responsible for organizing behavioral states). The second part of the analysis will introduce a first-mover model to understand the sequential behaviors of individual neuronal cell identities that underlie connectogenesis and the establishment of behaviors in the larval and adult worm.

### Large-scale Morphogenetic Trends

To momentarily step back from issues of terminally differentiated cell identity, we consider the birth time of the developmental precursors of connectome cells (Sulston et.al, 1983). Considering bursty events that precede the emergence of neuronal cells might provide information about large-scale morphogenetic trends at the organismal level. Figure 3 shows a complex distribution for the birth of developmental cells across the so-called first proliferation phase (Altun and Hall, 2011). While the birth of developmental cells occurs in aperiodic bursts during this interval, there are three bursts of particular interest to our sampling of neuronal cell births. The first burst of births are shown in red, and occur from 210 to 285 minutes. Next is a burst of developmental cell births shown in orange, and occurs from 285 to 345 minutes. While there are no developmental cell births from 345 to 365 minutes, there is a third burst of births from 365-400 minutes of embryogenesis. Based on a prior analysis of the data (Alicea et.al, 2019), bursts of developmental cell births may precede rounds of neurogenesis. The aggregate measure shown in Figure 3 is consistent with the finding that cellular specialization is coordinated with rounds of cell division, although this does not result from a specific cell division-related mechanism (Alberts et.al, 2002). The first few neurons and major expansion in the number of neurons occur immediately after the first two bursts of developmental cell proliferation.

**Figure 3.**
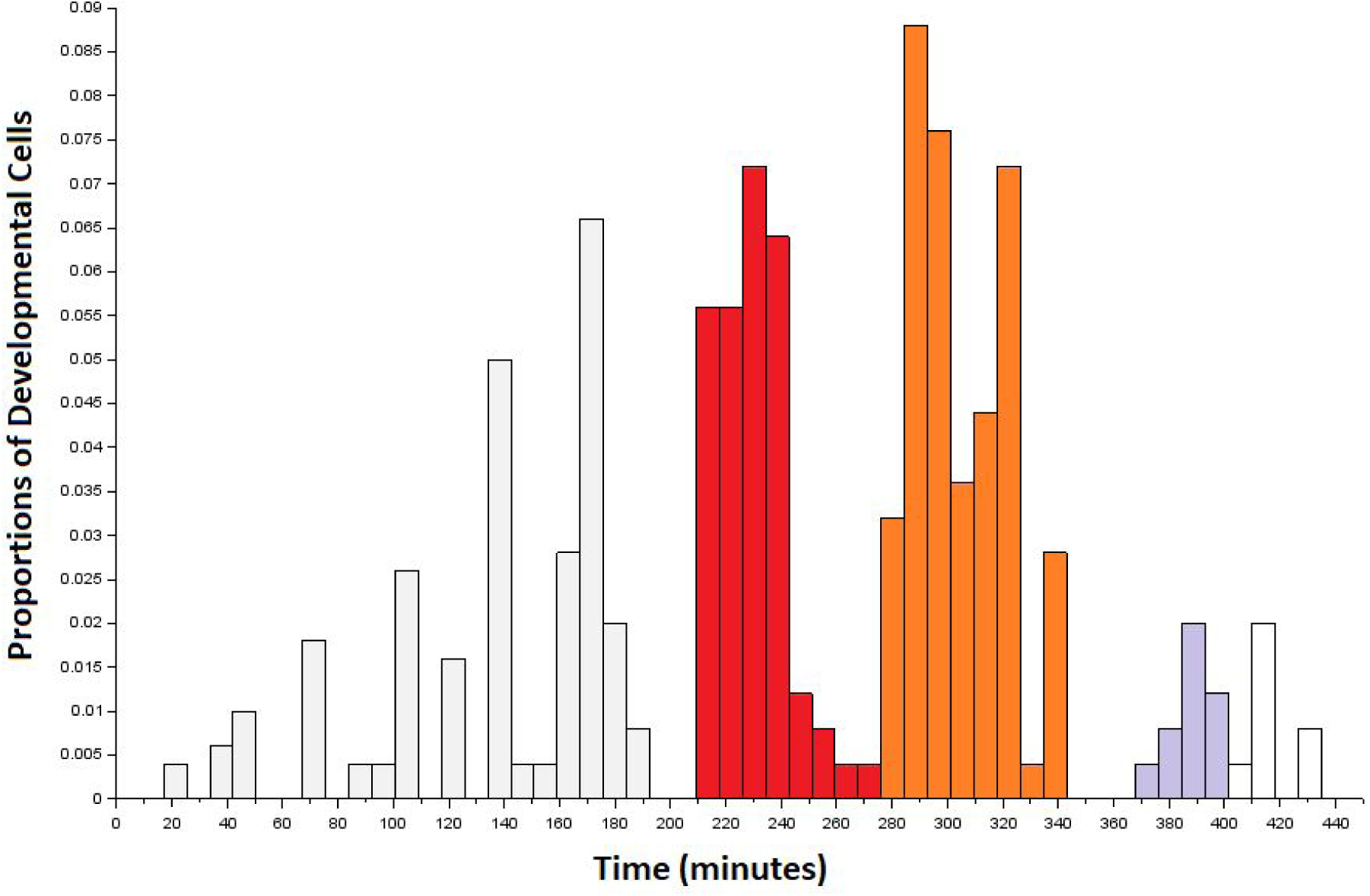
Developmental precursors to the first connectome cells to terminally differentiate in the embryo. Data are arranged as a histogram with bins of size 8. Colored bins correspond to the first (red), second (orange), and third (gray) bursts of developmental cells that overlap with the terminal differentiation of neurons from 265 to 400 minutes post-fertilization. The first burst of developmental cell birth (red) precedes our sampling window, but may contribute to neurogenesis during this time. The third burst (gray) is shaded up to the end of the sampling window.

Connectogenesis from 265 to 400 minutes post-fertilization. At 265 minutes, we have one cell that emerges: RMEV. RMEV is a ring motor neuron, and is part of the RME class. As this is positioned in the middle of the connectome anatomy, it is a symmetric single cell. According to Gendrel et.al (2016), RMEV forms part of the GABAergic nervous system in *C. elegans*, which constitutes nearly half of the adult *C. elegans* connectome. While RMEV serves as a founder cell (first cell to emerge in connectogenesis) for this network, its functional role as a core node in this subnetwork is not clear.

For the 280 minute connectome, there are five terminally differentiated neurons: RMEV (born at 265 minutes), and two symmetric pairs of neurons (ADFL/R and AWBL/R). ADFL and ADFR have been identified as amphid neurons that detect volatile compounds (Bargmann, 2006). Looking back at Figure 1, this represents the emergence of components in a functional circuit well before there are any environmental or activity-dependent signals to shape their fate. As harbingers of neuronal function, AWBL and AWBL are also important for their role in the NSY-5 network. About half of these cells share the same lineage, and the NSY-5 network as a whole is required for stochastic, asymmetric gene expression (Alqadah et.al, 2016).

The 290 minute connectome expands to 38 terminally differentiated neurons distributed outside of what will become the head for the first time (see Figure 4). There are a number of neuronal pairs born at this time, as well as several asymmetric terminal differentiation events. We can focus on two groups (AV and IL2) to appreciate the beginnings of functional architecture established at this point in development. Consider AV neurons born at 290 minutes of development (AVDL, AVDR, AVJL, AVJR, AVKL, AVKR, AVL). While we cannot confirm a direct functional role for these cells with WormBase, AVD neurons serve to mediate behaviors such as backwards escape following an anterior touch stimulus. Meanwhile, IL2 neurons (IL2DL, IL2DR, IL2L, IL2R, IL2VL, IL2VR) begin to form the inner labial structure of the head and ultimately play a functional role in larval plasticity. IL2 plasticity is crucial for enhanced sensory capabilities during Dauer formation (Procko et.al, 2011; Androwski et.al, 2017). IL2 also expresses behaviorally-relevant proteins such as OSM-9 and EGL-2. IL2 neurons originate from the ABal and ABar sublineages (Wu et.al, 2011). In terms of connectogenetic function, IL2 has been implicated in nictation and dispersal behaviors during the Dauer stage (Albert and Riddle, 1983). These two examples suggest that entire groups of neurons are born at the same time, perhaps revealing developmental modularity (Lacquaniti et.al, 2013) that is partially, but not entirely, based on later behavioral function.

**Figure 4.**
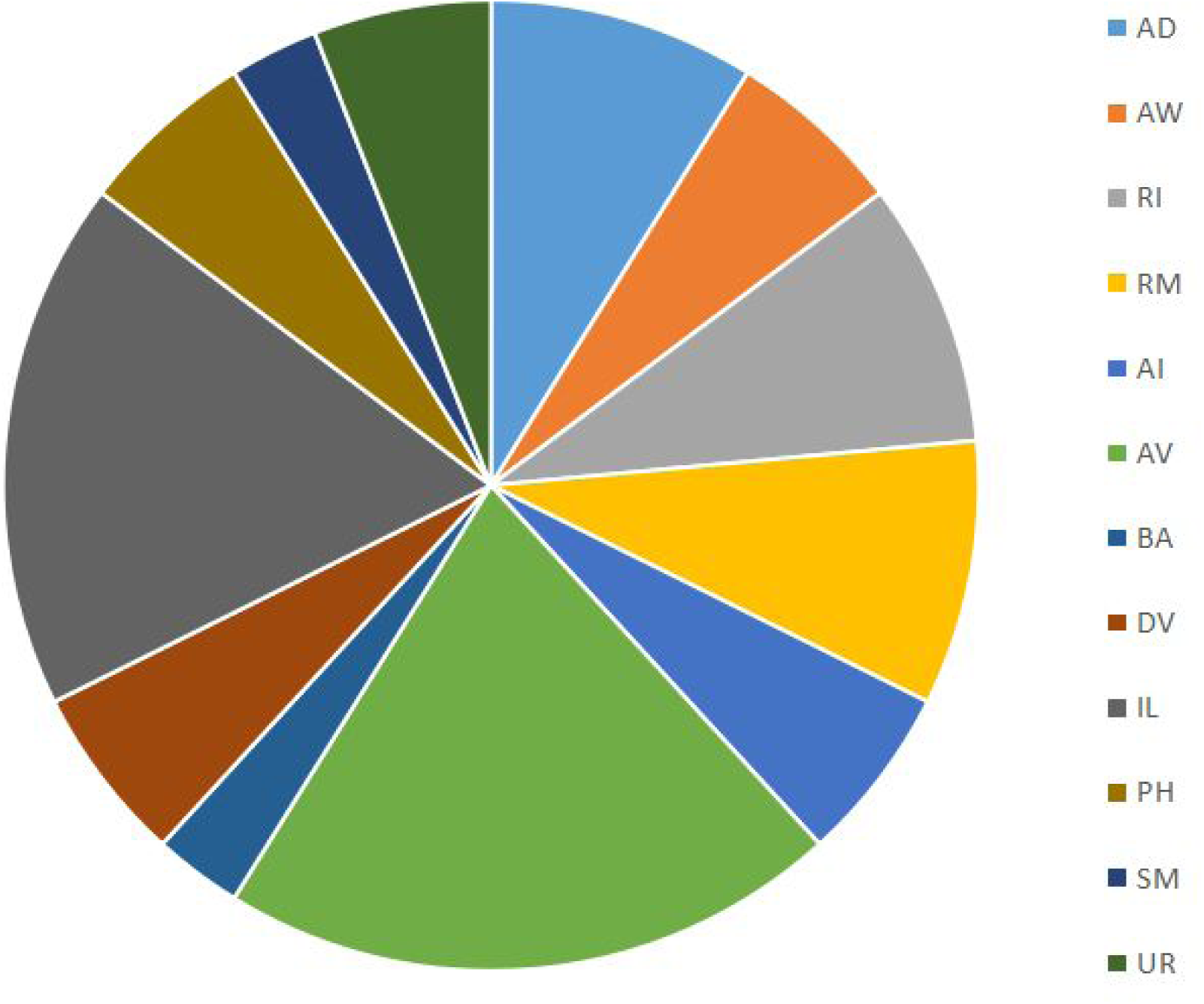
Distribution of cell types (based on families in Alicea et.al, 2019) born at 290 minutes. Cell families are nomenclature identities which share the same prefix and enable further analysis via mathematical (e.g. category-theoretic) techniques.

One cell pair that emerges at 300 minutes post-fertilization is AWCL/R. These are the Amphid Wing C neurons that will be essential post-hatch development for olfaction and form the basis for the NSY-5 subnetwork. In adulthood, AWCL/R neurons are unique among *C. elegans* olfactory receptors as they exhibit a fluctuation of intracellular calcium ions consistent with odorant stimulation, particularly the presentation and removal of the stimulus (Chalasani et.al, 2007). A computational model has demonstrated that the AWC cell pair are susceptible to noise, even when ion channels that mediate Ca^2+^ influx were included (Usuyama et.al, 2012). In terms of laying the foundation for a functional circuit, AWCL/R olfactory neurons work cooperatively with AIBL/R and AIYL/R interneurons to form an adaptive unit responding to food and odorants (Chalasani et.al, 2007). From our terminal differentiation temporal data, we can see that while AIBL/R and AWCL/R are born at 300 minutes, the AIYL/R interneuron does not appear until our 400 minute sampling interval.

A total of 87 new neurons differentiate between the 300 and 400 minute sampling intervals. Table 3 uses the concept of cell families (or theoretically useful groupings based on nomenclature identity) to show all new cells that occur in the 400 minute sampling interval that are not present in the 300 minutes sampling interval. The most abundant newly-differentiated neuronal families include members of the AV family (14 cells), RI family (9 cells), RM family (7 cells), and UR family (6 cells). Our 400 minute sampling interval also reveals the diversity of timing among inner labial (IL) neurons. For example, IL2 neurons are born at 290 minutes, whereas IL1 neurons appear at 400 minutes of embryogenesis. Despite this, each pair of neurons are cholinergic polymodal neurons (Pereira et.al, 2015) that have multiple mechanosensory functions in the adult (Goodman, 2006). While WormBase confirms that IL1 neurons enable mechanosensation in the adult worm, mechanosensation itself only becomes important later in development (post-hatch). As mentioned during the 280 minute analysis, we see neurons that form a functional circuit emerge well before the associated behaviors themselves.

**Table 3.**
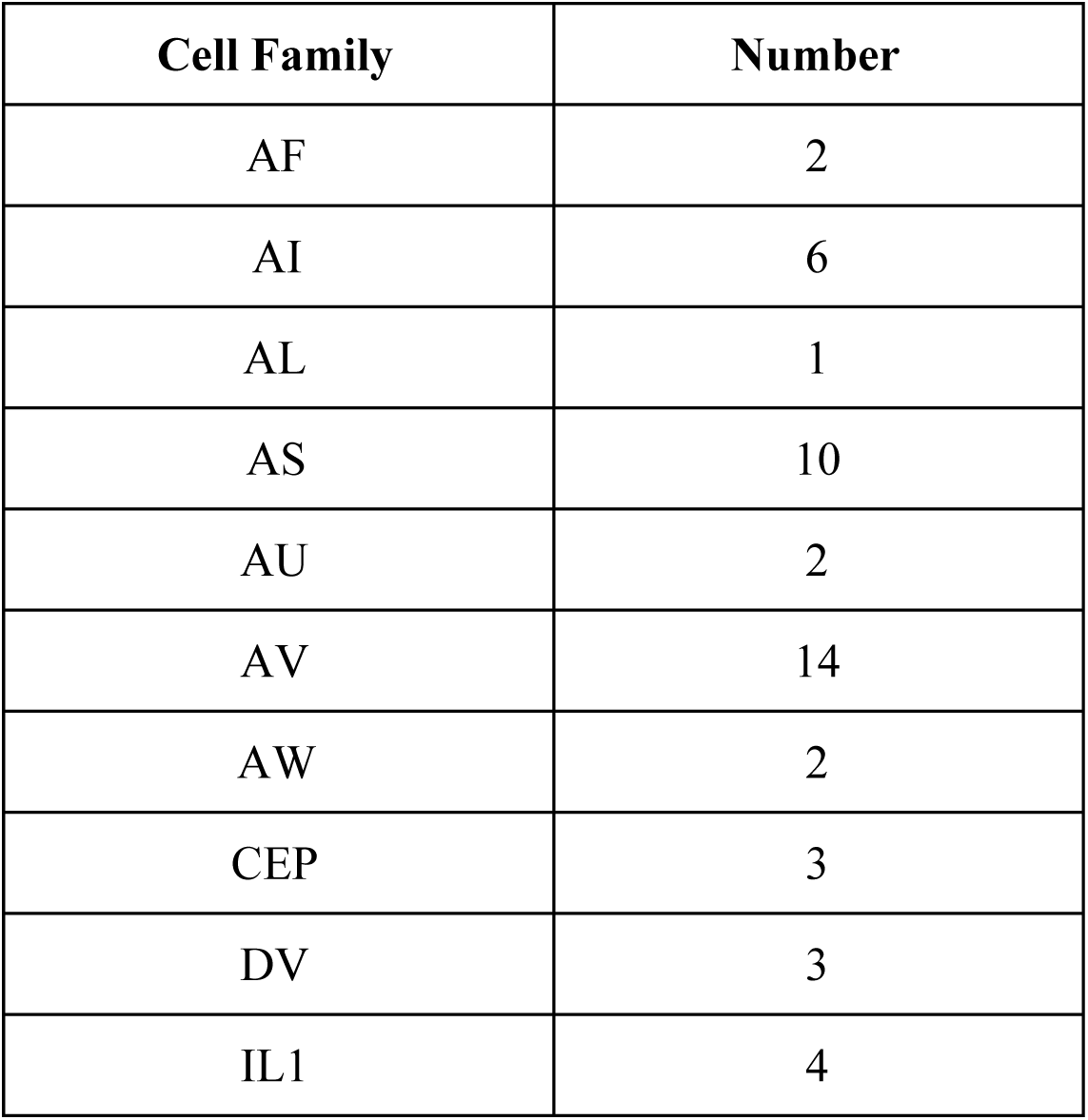
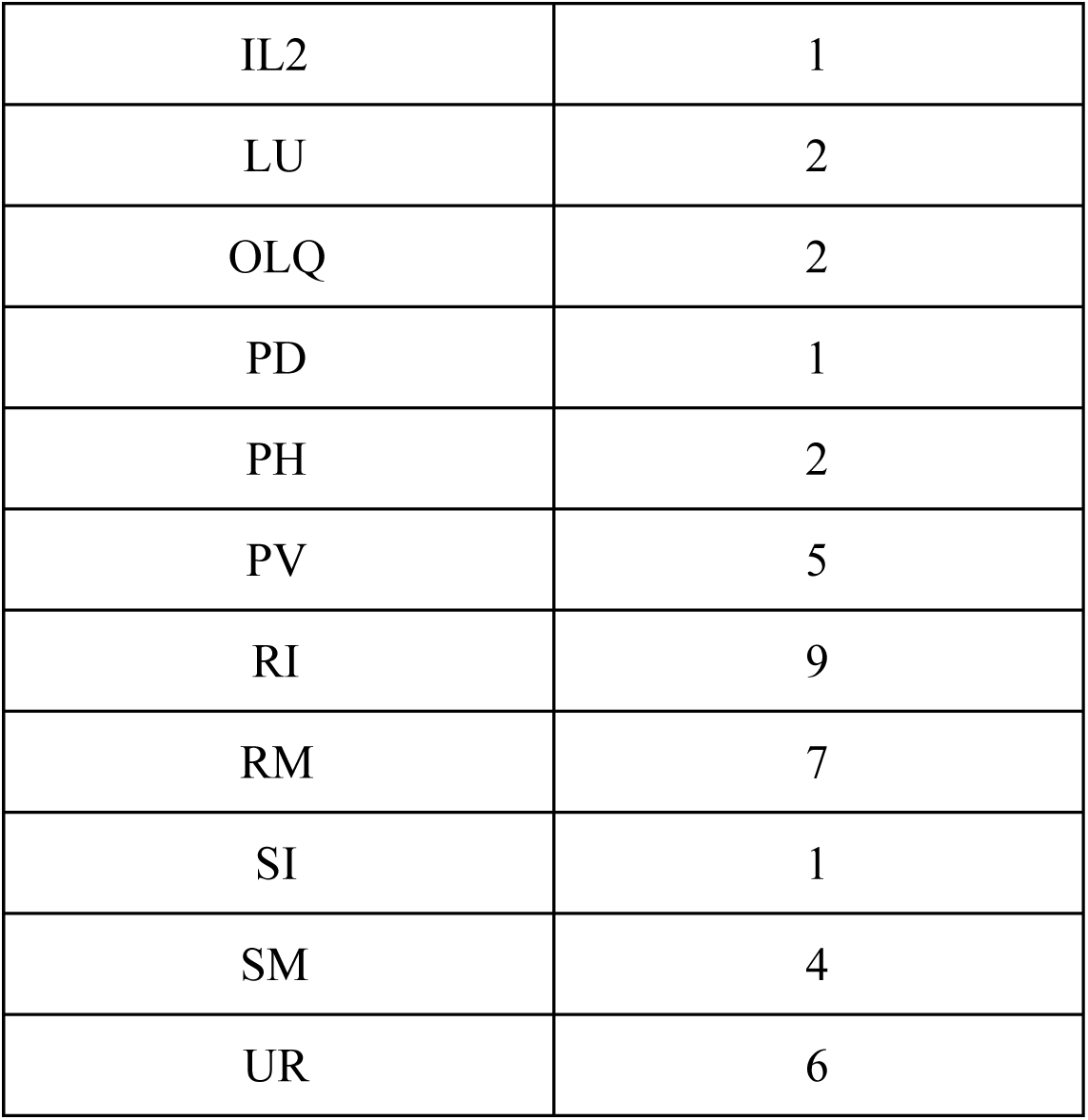
Terminally differentiated neurons present at 400 minutes of development versus 300 minutes of development. Cell family classifications are used, and a distribution across these families is provided.

| Cell Family | Number |
| --- | --- |
| AF | 2 |
| AI | 6 |
| AL | 1 |
| AS | 10 |
| AU | 2 |
| AV | 14 |
| AW | 2 |
| CEP | 3 |
| DV | 3 |
| IL1 | 4 |
| IL2 | 1 |
| LU | 2 |
| OLQ | 2 |
| PD | 1 |
| PH | 2 |
| PV | 5 |
| RI | 9 |
| RM | 7 |
| SI | 1 |
| SM | 4 |
| UR | 6 |

Connectogenesis post-400 minutes. There are a number of organismal level developmental events that occur after the 400-minute mark that are relevant to neuronal function. The 400 minute mark defines the beginning of the comma stage of development, where the head becomes thicker than the body and elongates correspondingly (Christensen et.al, 2015). Approximately 450 minutes post-fertilization (see Figure 1) marks the first evidence of axonal outgrowth (Morck et.al, 2003; Taylor et.al, 2010). From this time until hatching of the egg (800 minutes; Sulston et.al, 1983), we observe the onset of asymmetric gene expression in symmetric pairs of neurons. This is exemplified in neurons constituting the NSY-5 network (Hobert et.al, 2002; Taylor et.al, 2010). There is also an increasing asymmetry of phenotype outside of the connectome, as the hypodermis, muscle syncytium, and intestines all begin to take shape. This period between the embryonic neuronal birth and the onset of connectome-related behavior is an interesting period of time which will be discussed later in the paper.

Timing of NSY-5 subnetwork. Due to a number of functional attributes, the innexin-dependent NSY-5 network is worth examining in more detail. The NSY-5 network exhibits stochastic differentiation (Johnston and Desplan, 2008), which decouples the identity of developmental lineages from the transcriptional activity of terminally differentiated cells (Packer et.al, 2019). The first neurons in this network appear at 280 minutes, and continue to appear until after 400 minutes post-fertilization. At least one set of neurons in this network (AWCL/R, or Amphid Wing C cells) are influenced by calcium influx through voltage-gated calcium channels (Hseih et.al 2014). In particular, the NSY-5 network is required for left/right gene expression asymmetry in AWC neurons.

Overall, this network consists mainly of cells associated with Amphid Sheath and Neuropil, or function as Interneurons and Sensory Neurons. Left-right asymmetry of cell differentiation accounts for about 1/3rd of all terminally differentiated cells in *C. elegans*. Among AWC neurons, functional asymmetry is much more common than birth asymmetry. Birth asymmetry is greater than 50 minutes in only two pairs of NSY-5 cells (ASHL/R and PVQL/R). By comparison, symmetric induction of NSY-5 in AWC neurons occurs 150 minutes after their birth time, which is well after birth time but still falls within pre-hatch development (Taylor et.al, 2010). Beyond consideration of birth timing, there are only two neuron pairs (out of 98 total) for which we understand the underlying basis for their asymmetry: AWCL/R and ASEL/R. While many more of the 96 other pairs of neurons exhibit genetic and/or functional asymmetry, information regarding their function is currently lacking (Hammarlund et al. 2018).

Critical neurons and their cholinergic expression. To conclude with this survey, we will turn to the intersection of network functionality and the role of cholinergic neurons. Through a series of computational experiments, Kim et.al (2016) have identified 12 neurons that are critical for connectome function and involve 29 critical synapses. Critical neurons are highly connected compared with other neurons in the same network, and serve to bridge between different modules and reduce the overall number of connections across the connectome. Such neurons are central to a connectome and must be present in order for the various subnetworks of an adult connectome to remain functional.

In the face of environmental challenges and mutation, these cells enable the connectome to remain resilient (Gao et.al, 2016) without evolving new innovations. Nine of these 12 critical neurons are born in the hermaphrodite connectome prior to the initiation of the twitching phase (470 minutes). Six of these nine cells have cholinergic function: AVAL/R, PVCL/R, PVPR, and AVER. All of these cells are positive for *unc-17/cha-1* and *cho-1* expression (Pereira et.al, 2015). These cells also express *C. elegans* homologues of ACE genes (Schaefer, 2005), which are associated with a host of diverse functions.

Towlson et.al (2015) have identified a core set of rich club neurons, which also serve as critical for connectome function. This set of 11 cells differs slightly from the Kim et.al (2016) list, and also includes cholinergic neurons AVBL/R, AVDL/R, and AVEL. The map of Pereira et.al (2015) shows that not only are these cells ventral cord interneurons with cholinergic function, they also receive ACh inputs from other cells. It is not clear whether there are additional molecular and/or electrophysiological properties that differentiate rich club neurons from other neurons in the connectome in an *a priori* manner, but hierarchical relationships with other neurons in the connectome can be approximated using a first-mover model.

First-mover analysis with synaptogenesis data. During the course of connectogenesis, networks grow in terms of their size (cells represented as nodes and connections represented as edges) over time. These growing and actively-connecting structures are called *new world networks*, and require a means to integrate new nodes into the existing network (often defined as a *model of competitive network growth*). The model of competitive network growth predicts that new nodes compete to connect with existing cells in the network, while existing nodes compete to gain new cellular partners while also retaining old cellular partners. This results in *context-specific models of network attachment*, defined as a model of competitive network growth that occurs in a specific organism, genetic background, or environmental setting. With our first-mover model, we assume that Stackleberg competition is a model of network growth. It is also assumed that the structural connection rules of Pathak (2020) and the synaptic relationships proposed by Witvliet et.al (2020) serve as the context-specific model of network attachment.

In Figure 5, we integrate the synaptic formation data with birth time data for each cell. A simple comparison of the difference in birth time yields a fairly regular distribution of points around zero difference, suggesting that individual events are not informative of strategies employed by cells given a wider range of contexts. Interestingly, the distribution for developmentally-specific and stable synaptic relationships are skewed slightly to presynaptic cells being born *later* than postsynaptic cells.

**Figure 5.**
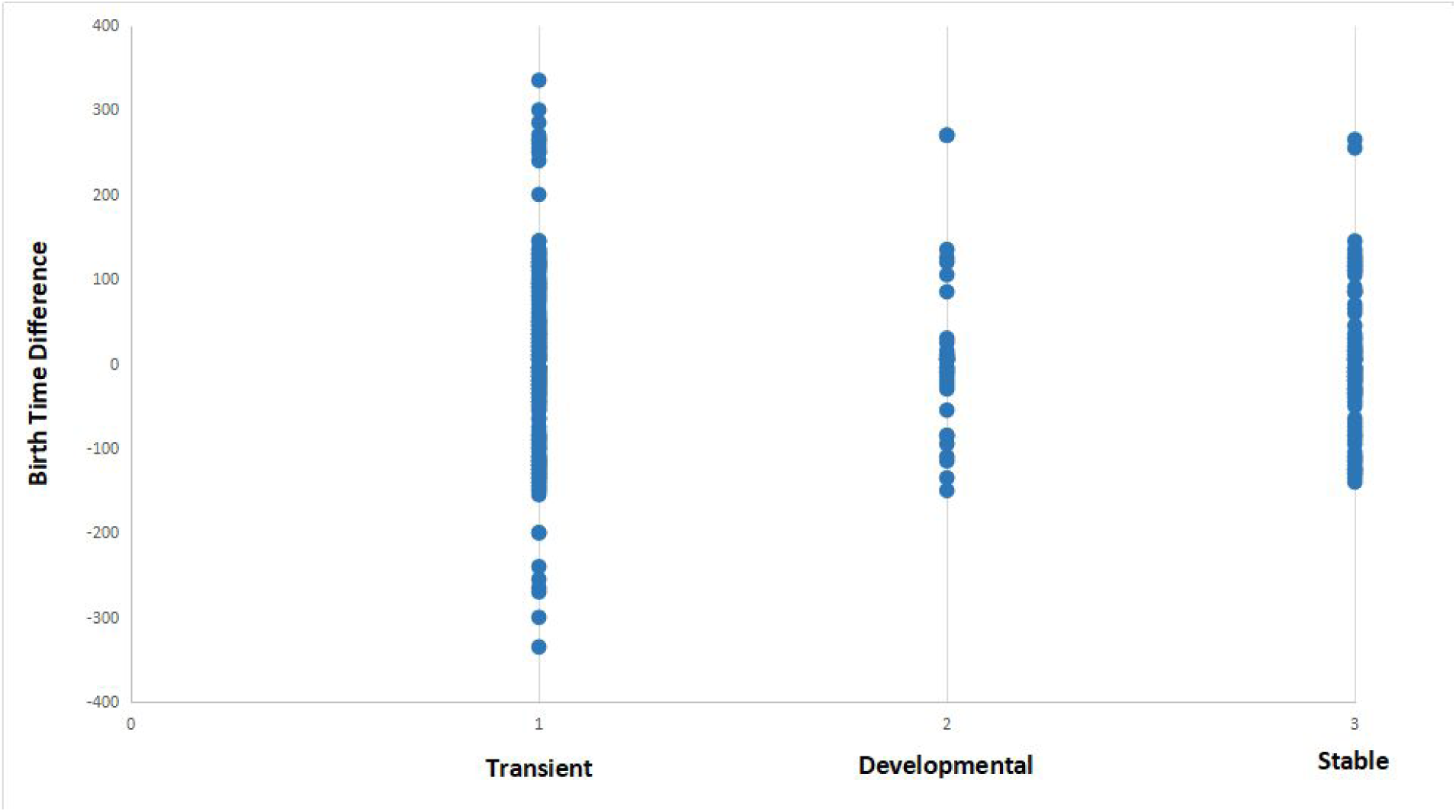
Distribution of synaptic relationships (for three different functional states) in terms of the difference in birth time between pre- and postsynaptic cells. See Methods for description of transient, developmental, and stable classifications. Negative values represent relationships where the presynaptic cell is born first in embryogenesis, while positive values represent relationships where presynaptic cells are born after the postsynaptic partner. Transient synapses exhibit an almost symmetrical distribution of positive and negative birth time differences. When compared to developmental and stable synapses, transient synapses include all connections that involve a strongly negative birth time difference (greater than 100 minutes). This suggests that connections resulting from negative birth time differences are more selective for functionally specific types of connections.

First-mover dynamics are based on scenarios where all players (in this case cells) have variable access to a global informational state (state of every neuron in the worm). There are five types of strategy based on the birth time of the observed pre- and postsynaptic couplings: negative-positive (N-P) coupling, exclusive OR (XOR) first-mover, exclusive OR (XOR) second-mover, exclusive NOR (XNOR), and positive-negative (P-N) coupling. The prevalence and outcomes of all strategies extracted from the data are shown in Table 4. To make these data relevant to formal strategies, we make two assumptions about how connection behaviors exhibited by cells can be viewed as decision-making. Cells make their decision about which strategy to employ not based on individual contacts, but on a memory of prior states.

**Table 4.** Frequency of different first-mover strategies among cells in developing *C. elegans* connectome.

|  | Frequency of strategies in population | Proportion of synaptic types for cells that exhibit a particular strategy |  |  |
| --- | --- | --- | --- | --- |
|  |  | Transient | Developmental | Stable |
| **N-P coupling** | 41.5% | 38% | 36% | 43% |
| **XOR First Mover** | 7.5%* | 8% | 24% | 6% |
| **XOR Second Mover** | 18.9%* | 8% | 24% | 8% |
| **XNOR** | 15.1%* | 17% | 24% | 13% |
| **P-N coupling** | 45.3% | 86.2% | 91.1% | 85% |

\* overlap with N-P coupling category.

Aside from identifying specific strategies, we can also identify cells that employ these strategies. The N-P strategy is exhibited by the following cells: ADF, ASI, AUA, AVA, AVB, AVD, AVE, DVA, DVC, IL2D, IL2L/R, IL2V, PVP, PVR, RIA, RIB, RIG, RIH, RIR, URB, URYD, and URYV. It is of note that the N-P strategy overlaps with the XOR and XNOR strategies. Cells exhibiting the XOR First-mover strategy include: ALM, BDU, DVC, and IL2D. The XOR Second-mover strategy is exhibited by the following cells: AVA, AVB, AVD, AVE, AWB, BAG, IL2L/R, RIA, URYD, and URYV. Cells exhibiting the XNOR strategy include: ADF, ASI, DVA, IL2V, PVR, PVT, SAAD, and SAAV.

In terms of overlap, the N-P and XNOR strategies are co-employed for 5 out of 8 possible cells, while overlap between N-P/XOR First-mover and N-P/XOR Second-mover are co-employed for 2 out of 4 and 8 out of 10 possible cells, respectively. The fifth strategy (P-N) is notable for no overlap with any of the other four strategies. Cells exhibiting the P-N strategy include: ADA, ADL, AFD, AIA, AIB, AIN, AIY, AIZ, ASE, ASG, ASH, ASJ, ASK, AWA, AWC, FLP, OLL, OLQD, OLQV, PVC, RIF, RIM, RIP, and URX.

To reveal this, a raw averaging of birth time difference is not informative due to the great variety and sometimes trivial difference in birth times between cells. The consequences and broader implications of each strategy on the synapse connectomic data are shown in Figure 6, which unpacks the connectivity-birth time order relationship that is not obvious from the analysis shown in Figure 5. As a measure of connectivity, average synaptic density is used to understand whether or not the cells born early are contributing disproportionately to the pool of synapses observed across development.

**Figure 6.**
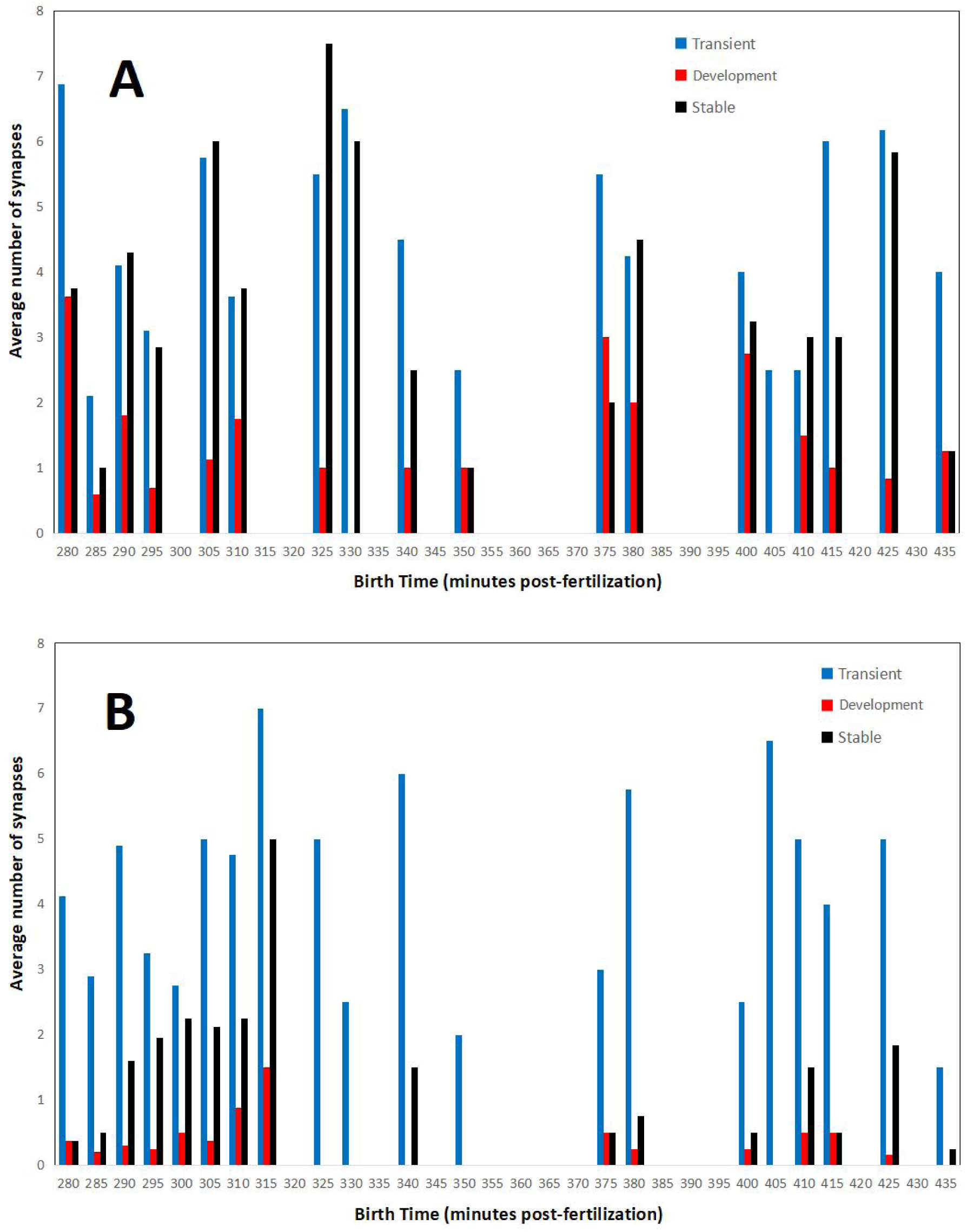
Average number of presynaptic (A) and postsynaptic (B) connections for cells born at a specific time in embryogenesis. Synaptic counts for each category and time point are normalized by the total number of cells born at each time point. Bars in order for each time point: transient (left, blue), developmental (middle, red), and stable (right, black).

There are two distinct patterns that emerge when comparing pre- and postsynaptic connections. For presynaptic connections, there appears to be a cyclic pattern over time. Meanwhile, postsynaptic connections consist primarily of neurons born both earlier and later in embryogenesis. Presynaptic connections of all classifications (transient, developmental, and stable) are made by cells born in rounds of 10-15 minutes each. There are also times for which no presynaptic connections are associated. This may even be related to the cyclical nature of developmental cell birth shown in Figure 3. For postsynaptic connections, a majority of the developmental and stable synapses are born early (from 280-315 minutes). As a result of postembryonic plasticity, transient connections are numerous and associated with cells born at all times. These patterns might also serve as an indicator that the strategies presented here operate across embryogenesis, and that terms such as first and second mover are relative to each cell’s local context.

At this point, we might ask why cells select a specific strategy in the first place. One heuristic might be that cells have a way of detecting the birth time of their fellow cells. This might be suitable for the *C. elegans* example, where cells exhibit programmed differentiation and the connectome size is relatively small. Yet since axonogenesis is delayed until most of the neurons have terminally differentiated, it is also not possible that cells connect to whoever is around them at a specific time. Another heuristic might involve sensing the difference in time since terminal differentiation between themselves and a potential synaptic partner. This is the logic behind the first-mover analysis, but it is still not clear how cell age maps to our defined strategies.

Figure 7 shows the cumulative presence of cells over time identified by their strategy. In this analysis, we find that cells exhibiting the N-P strategy are born on average earlier than P-N strategy cells. In fact, cell birth times for P-N strategy constituents lag behind those for all other strategies. This is perhaps the opposite of what is expected if cells detect birth time rather than a difference in age. However, recall that the first-mover model is a multi-step process in which a single cell can serve as a presynaptic partner to many cells simultaneously. In addition, the *C. elegans* connectome is characterized by densely-connected hubs which allows for few presynaptic partners to have many postsynaptic partners. On the other hand, cells exhibiting the XNOR strategy (the most refined first-mover strategy) are consistently born before cells exhibiting other strategies. XOR First-mover cells follow a similar pattern of early birth times, and it may be that strategies that explicitly exclude presynaptic partners born later than the cell in question are favored among cells that are born early in connectogenesis.

**Figure 7.**
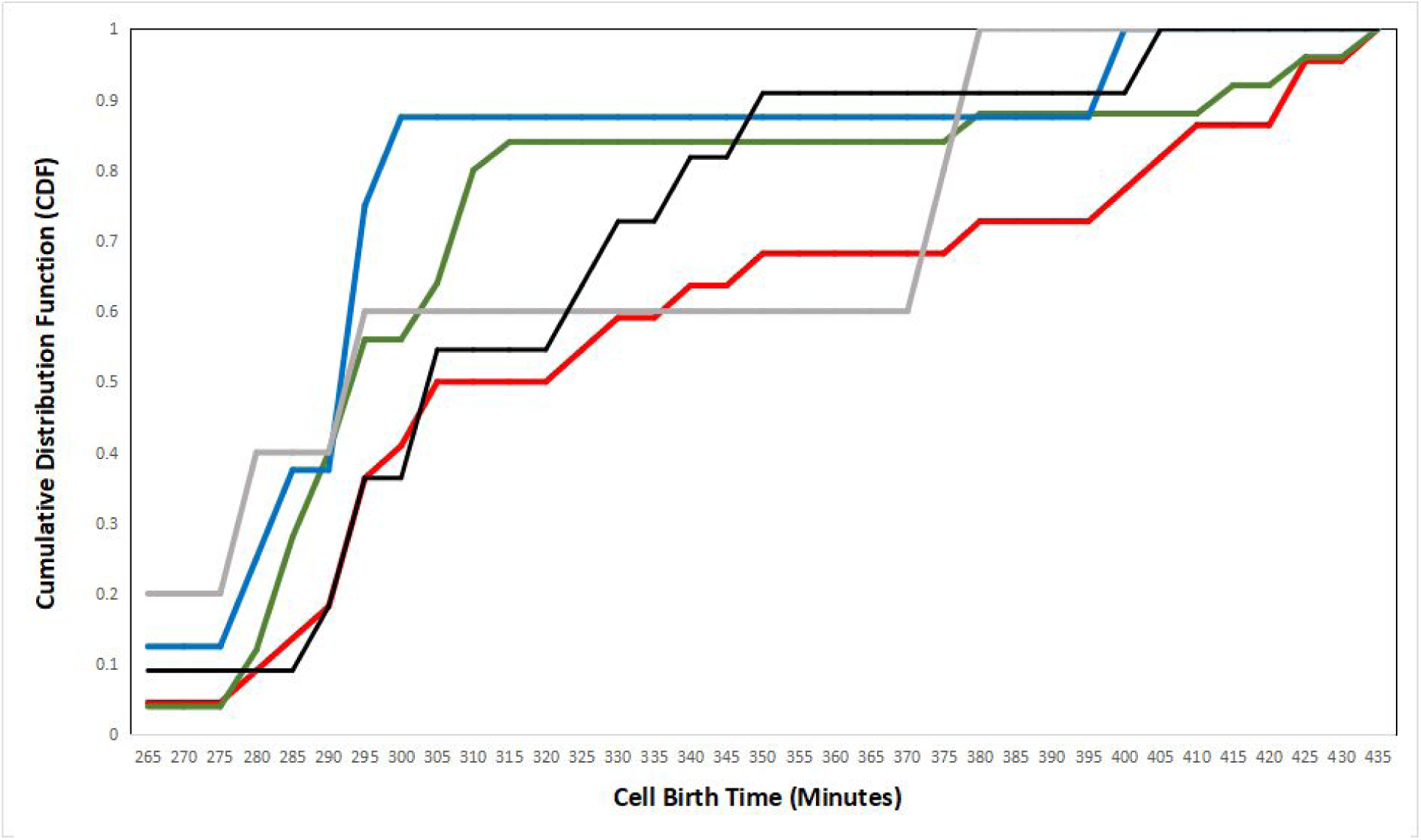
A plot of the cumulative distribution functions (CDFs) for each strategy, plotted as the birth time distribution of cells exhibiting single strategies. N-P is shown in red, XOR First is shown in gray, XOR Second is shown in black, XNOR is shown in blue, and P-N is shown in green.

There are a couple of insights to be gained from the first-mover analysis. The first insight involves the exclusive nature of first-mover strategy. While our analysis identifies tendencies of strategic behavior, two strategies (XOR first-mover and XOR second-mover) are pure strategies.

Of these two, XOR first-mover is the rarer strategy, suggesting one of two possibilities. It might be that the payoff is lower for presynaptic cells to establish connections with cells originating at a later embryonic birth time. Alternately (or perhaps concurrently), the XOR first-mover strategy might have a higher payoff for certain cell identities.

The second insight is that there are interesting properties for presynaptic cells that seem to determine their connection partners by relative developmental birth time. For presynaptic cells that are born after their postsynaptic partners, they dominate the number of synapses established in the population, despite constituting a little less than half of the total population. Meanwhile, presynaptic cells that are born before their postsynaptic partners demonstrate a sparsity of synaptic connectivity, almost as if there is a selective mechanism that limits the proliferation of synaptogenesis overall and favors developmental and stable synaptic connections over transients.

### Expansive Networks (Systems)

In cases of defined mutants, regulative development, or transcriptional noise/error, the first-mover game is played with imperfect information. When we are unsure about the identity or timing of cell differentiation, it is harder to predict the outcome of such a game. What is missing from the literature are ways to understand the expansion of developing neural systems. The fundamental idea of networks expanding according to the need of a given system (Bell et.al, 2017) is typically characterized in terms of connection principles such as propinquity (Gallos et.al, 2019) that apply regardless of whether a network is sociotechnical or biological. The first-mover model demonstrated here provides a different means of characterizing the expansion of network connectivity.

Importantly, multiple factors can be used to explain the often complex behaviors of connectome integration. Using microscopy data that shows budding axons in development, Pathak et.al (2020) propose that these factors might include: birth time, developmental lineage proximity, fellow sisters cells, cells with a common fate, and bilateral pairs. Some of these factors indeed constrain connectivity to functionally-relevant partners, but also enforce features such as functional asymmetry that are observed in adult worms.

Yet why is a first-mover model superior to simply relying on a set of correlations? One important reason involves the discovery of assembly rules for an emerging connectome. The formation of a connectome as a network that expands in size has been characterized as so-called New World Networks (Hilgetag and Goulas, 2016). New World Networks not only utilize multiple connection rules simultaneously, in a way that allows for equilibrium shifts in the first-mover model as described in Stone et.al (2018). Secondarily, models of competitive network growth in a network provide context-specific models of network attachment (Szalkai et.al, 2017). These connection rules are described as intentional strategies, but ultimately rely upon the activity-dependence of ion channels and gene expression more generally.

## Discussion

This paper presents several key findings that flesh out aspects of the developmental connectome. We provide a guided tour of early developmental events related to newly-differentiated neurons and their physiological context. In addition, we systematically walk through the developmental connectome at different periods of embryogenesis and attach information regarding that provides context about what is observed in adult cells and how they fit into the connectome. The point of view presented here is tempered by the finding that gene expression varies widely across development, even in *C. elegans* (Hashimony et.al, 2015). This does not invalidate our attempts to trace the steps of connectogenesis and establish linkages between embryo and adult. It does, however, highlight how fluctuations at the molecular scale obfuscate (or contribute to) relative persistence at the anatomical scale. We then analyze the establishment of synaptogenesis during larval development using a theoretical model that ascribes strategies to each cell. Excluding influences due to post-embryonic connectome plasticity (Bhattacharya et.al, 2019), it is the agency of larval synaptogenesis that largely determine the establishment of goal-directed behaviors.

### Limitations of Approach

There are several limitations of this approach. The first limitation is the use of informatics as an inferential tool. While we have been able to integrate multiple sources of data, direct observations of the system itself are as of yet not available. Yet despite these limitations, we can provide a significant view of an emerging nervous system. This can be done in *C. elegans* for two reasons: a deterministic cell lineage (Sulston et.al, 1983) and eutelic adult phenotype (Sulston and Horvitz, 1977). Even in cases where there is variability in the developmental process across individuals, we should expect this variability to be predictable (Azevedo and Leroi, 2001).

Despite the limitations of such an approach, we can also use the information presented here to better understand the relationship between connectome formation and behavior. This transition is poorly understood from a systems point-of-view.The information provided in this paper will allow us to reconstruct networks in a number of ways. The first involves being able to expand or contract the number of cells and their gap junction connectivity present at early developmental time points. Coupled with information about tendencies in synaptic coupling, we can estimate the number and distribution of edges in an emerging connectome. This allows us to overcome the limitations of defining the connectome exclusively in terms of electrical (gap junction) or chemical (synaptic) connectivity (Morgan and Lichtman, 2013). While this might allow us to model how sensorimotor behaviors are enabled immediately before and after hatch, it also has an effect on understanding how routing strategies in the connectome might have their origins in early development.

### Relevance of Networks and First-mover Dynamics

This work also provides insights into the presence of biological large-world networks. While the adult *C. elegans* connectome has been characterized as a small-world network (Watts and Strogatz, 1998), this connectivity pattern might not have this same pattern in early development, when the number of cells, axonal connections, and synaptic connections are in the process of being formed. In many instances, such relationships have yet to be established. network expands faster than shortcuts can form. While theory credits forms of cumulative advantage (deSolla Price, 1965) such as preferential attachment, developmental analysis (Nicosia et.al, 2013) suggests that preferential attachment fails to account for biphasic growth in the number of cells across both embryonic and postembryonic development. Identification of emerging molecular signatures along with a formal model of developmental emergent complexity moves us closer towards biologically-based and mathematically-rigorous models of developmental connectomics.

The assignment of strategies to cells may seem presumptuous, but it does provide a new view of how the connectome emerges. One way to understand the impact of such strategies on consequential features of the adult organism is to evaluate strategies employed by the rich club of connectivity. The overlapping strategies N-P coupling and XOR Second-mover are preferred for 10 of the 14 rich club neurons. An additional rich club neuron exhibits N-P coupling and XNOR strategies. This suggests an important role for both birth order and developmental contingency in the formation of hierarchical and non-random structure in the connectome.

We can also connect the first-mover tendencies with the NSY-5 network. This is particularly relevant to connectome development for at least two reasons. The first is related to the timing of NSY-5 network formation: NSY-5 expression begins in mid-embryogenesis, and is strongest in late embryogenesis and the L1 larval stage (Chuang et.al, 2007). Secondly, Schumacher et.al (2012) suggests that functional asymmetries in the NSY-5 network are based on electrical connectivity. The extension of these asymmetries to synaptic connectivity might be reflected in the strategies employed by the 28 cells in this network, as 16 of them (ADA, AIY, AIZ, ASK, ADL, AFD, ASH, and AWC) exhibit the P-N strategy, with the rest of the cells exhibiting various N-P strategies. With roughly half of the cells in this network choosing between one of two strategies to form chemical connections, this could be interpreted as cells setting up two different functional modules within the same network.

### Broader Implications

There are several broader implications to this work. The first is to flesh out a model of explicit steps in relation to the development process and with respect to an emerging connectome. The second involves informing developmental models of nervous system function. This provides a basis for understanding the emergence of specific behaviors and behavioral circuits. First-mover models can contribute to the study of development by characterizing the role of developmental contingency, or the order in which cellular functional and structural configurations become manifest in the phenotype. The presentation of information here is a first-step (and in some cases significantly more) towards both parameterizing models of embryogenesis and a unified understanding of the first steps towards autonomous behavior.

As the *C. elegans* connectome is considered analytically tractable, we hope that the extraction and analysis of neuronal populations at different points in embryonic development can be applied to other small connectomes (<1000 neurons). The polychaete (Windoffer & Westheide, 1988; Hochberg, 2009) and the larval tunicate *Ciona intestinalis* (Bezares-Calderon & Jekely, 2016; Ryan et.al, 2016) are two such examples. The tunicate model is especially interesting, as it provides an opportunity to study asymmetry. Yet we also see how the emergence of behaviors are context-specific, as swimming in *C. intestinalis* (Zega et.al, 2006) emerges in a way that is fundamentally different from behaviors such as pharyngeal pumping in *C. elegans*.

Development and Neuronal Criticality. Considering neuronal criticality more broadly, Towlson et.al (2013) have defined rich club neurons in the adult phenotype as highly-connected neurons and serving a special functional role due to their ability to connect different modules. One question for future research is whether or not the rich club originates from the rules of preferential attachment (Barabasi & Albert, 1999), or if their functional attributes make them prone to being highly-connected. As they tend to be born between 290 and 350 minutes of development (Alicea, 2019), birth order is not the preferential criterion for this functional role. In any case, we can observe the signature of a small-world network (Watts & Strogatz, 1998) in both the *C. elegans* and *Ciona intestinalis* larval connectomes (Bezares-Calderon & Jekely, 2016). This suggests whatever interactions are determining critical structure is common across developing small connectomes.

### From Embryogenesis to Behavior

Our ability to infer behavioral pathways as they form in embryogenesis can yield information about how and why behavioral circuits form in development. To see the value of this information, we can recall Tinbergen’s four questions with respect to an organismal trait: its structure, its function, its developmental origins, and its evolutionary origins (Bateson and Laland, 2013). In this paper, three of these (structure, function, and development) are addressed. This inquiry proceeds in a manner consistent with the aims of neuroethology in that behavior is best understood by focusing on a diversity of stereotypic behaviors (Hoyle, 1984). Based on this analysis of *C. elegans* data, an order of origins can be proposed: developmental processes (cellular differentiation) generate structure (the birth of circuit components), which in turn leads to function (connections between neurons and the generation of behaviors). To further support this hypothesis, structures such as muscle autonomously generate behaviors prior to innervation by axonal connections. We will now review such behavioral transitions to demonstrate how structure comes before function.

The first visible motor activity in the embryo is twitching (see Figure 1), which begins at 470 minutes of development (Kim et.al, 2016). Twitching is defined by spreading calcium waves in muscles and calcium transients in motor circuit neurons (Ardiel et.al, 2017). While the earliest twitches are purely myogenic in origin, later twitches (once synaptic connections are established) are controlled by the cells in what will be the adult motor circuit (Ardiel et.al, 2017). One of the first complex behaviors exhibited by *C. elegans* is pharyngeal pumping (see Figure 1). Pilon and Morck (2005) make two observations about the origins of the pharynx. One is that the morphological components of the functional pharynx (organ shape and position) become established between 430 and 760 minutes of development (Portereiko & Mango, 2001). The other is that rhythmic contractions of the pharynx can occur in the absence of neuronal control (Avery & Horvitz, 1989). Efficient pharyngeal pumping (which does require neuronal control) is vital to life outside of the egg, and thus occurs at around 760-780 minutes of development (Chisholm and Hardin, 2005). This type of structure-function relationship is observed in other small connectomes as well. In *C. intestinalis* larvae, both tail flicks and phototropisms are generated by muscles before there are connections with neurons (Bezares-Calderon & Jekely, 2016).

Throughout early larval development, we observe the emergence of behaviors in specific functional circuits related to escape responses and mechanosensation and locomotion more generally (Bozorgmehr et.al, 2013). In the case of locomotion, changes to both connectome size and synaptic connections to muscle groups occur during the early larval stages (White et.al, 1978). This affects refinements in movement behavior, particularly with respect to post-embryonic developmental plasticity. By the time the *C. elegans* egg hatches, we observe the beginnings of a network specialized for escape response behaviors. This escape response involves forward and backward movements with respect to touch stimuli (Maguire et.al, 2011). When a worm is touched on the anterior end, sensory cells ALM and AVM trigger a backwards movement by motor neurons AVD, AVA, VA, and DA (Pirri & Alkema, 2012). By contrast, touching a worm on the posterior end triggers sensation by cells PLM and PVM, which results in forward movement generated by PVC, AVB, VB, and DB motor neurons (Pirri & Alkema, 2012). Additional cells such as ASH and RIM are involved in the control of head and recoil movements, respectively (Campbell et.al, 2015; Zhao et.al, 2003). While our first-mover strategies do not explain the emergence of this circuit, cells associated with two behavioral components (reversal and head control and recoil) correspond to two sets of strategies: the P-N strategy for cells ASH and RIM head control and recoil), and the XOR first-mover and XOR second-mover strategies for cells ALM, AVA, AVB, and AVD (reversal).

### First-mover Model as the Basis for Simulation

There are a number of potential uses for first-mover models to bridge function and behavior. Clement (1987) suggests that a solid mapping between neurotransmitters, sensory devices, muscles, and organ systems allows us to gain knowledge with respect to function. We can map our first-mover model to neuroethological diagrams, which provides a means for establishing this mapping. This in turn helps to clarify some limitations of a purely informatics and modeling approach. While direct observation of the connectome is sparse, first-mover models allow us to postulate potential mechanisms for initiating the self-organization of developmental complexity.

This leads us to a question: can we simulate or even replicate the process of “booting up” a connectome? Using physical connectivity information (gap junctions) in tandem with time-series sampling, we observe a number of steps to the construction of a connectome. At 280 minutes for example, we observe both what will become reciprocally connected pairs of cells (ADFL/R and AWBL/R) and a disconnected cell (RMEV) whose network partners have yet to be born. Observations such as these resemble what Betzel et.al (2016) term a generative connectome. As the adult connectome exhibits a hierarchical structure where some neurons are more central to the network (demonstrated in terms of more dense connectivity) than other cells, we must also ask how this comes to be. It has previously been found that early-born neurons tend to be both more highly connected and exhibit more longer-range connections than later-born neurons (Varier and Kaiser, 2011). Yet neuronal birth time is not particularly informative of the details of later synaptic connectivity (Kratsios et.al, 2015). In this paper, we propose that criteria such as preferential attachment or fitness might play a role in shaping the adult nervous system’s connectivity patterns. In turn, a first-mover model allows us to characterize these different hypotheses, in addition to synthesizing what is known with respect to the molecular and electrophysiological properties of connecting cells.

Another area in which this study would be of value is in informing simulations studies. One example from *C. elegans* involves the c302 project (Gleeson et.al, 2018), which is an attempt to simulate various aspects of the adult nervous system. Such a model adapted for embryonic and larval development would help to close the empirical gaps of the current approach. Of particular interest is the platform’s ability to simulate subcircuits and how this might be used to recapitulate the developmental process. Applying c302 to a developmental context might mean simulating early-stage ion channel expression (Rutenberg et.al, 2002) and the activity of ion channels in the absence of synaptic connections. This might be done by hypothesizing which ion channels become functional when among the cells switched on at different times. Yet despite the efforts of OpenWorm, two pieces of information regarding the underpinnings of behavior are still missing: a functional connectivity map at the biochemical level, and a systems-level map of experience-dependent plasticity (based on Schaefer, 2005).

The advancement of work on Developmental Braitenberg Vehicles (BVs - Braitenberg, 1984) might also allow us to evaluate the emergence of cells and activity patterns in the context of embodied development (Dvoretskii et.al, 2020). This takes the first-mover model a step further to provide both environmental and sensorimotor feedback, which play a role in bridging the gap between self-generated twitching and coherent sensorimotor behaviors (Narayanan & Ghazanfar, 2014). While Developmental BVs provide a connectionist view of an emerging connectome, these connections can be modulated by mechanisms that are activated or fluctuate during the process of interacting with its environment. Developmental BVs also rely on systems-level regulatory mechanisms such as fitness functions or Hebbian learning mechanisms. A similar type of mechanism is likely to be critical for developing biological connectomes as well. Brought into the context of electrophysiological function, we can look toward multilevel Developmental BVs that model gene expression using artificial Genetic Regulatory Network (GRN) models (Cussat-Blanc et.al, 2019) as a means to augment the function of a connectionist network.

